# Metabolomic Phenotypes Reflect Patient Sex and Injury Status: A Cross-Sectional Analysis of Human Synovial Fluid

**DOI:** 10.1101/2023.02.03.527040

**Authors:** Hope D. Welhaven, Avery H. Welfley, Prayag Pershad, James Satalich, Robert O’Connell, Brian Bothner, Alexander R. Vap, Ronald K. June

**Author notes:** <u>Corresponding Author</u> Ronald K. June, Ph.D. Dept. of Mechanical & Industrial Engineering, Montana State University, PO Box 173800, Bozeman, MT 59717-3800.

## Abstract

1.

**Background:** Post-traumatic osteoarthritis (PTOA) is caused by knee injuries like anterior cruciate ligament (ACL) injuries. Often, ACL injuries are accompanied by damage to other tissues and structures within the knee including the meniscus. Both are known to cause PTOA but underlying cellular mechanisms driving disease remain unknown. Aside from injury, patient sex is a prevalent risk factor associated with PTOA.

**Hypothesis:** Metabolic phenotypes of synovial fluid that differ by knee injury pathology and participant sex will be distinct from each other.

**Study Design:** A cross-sectional study.

**Methods:** Synovial fluid from n=33 knee arthroscopy patients between 18 and 70 years with no prior knee injuries was obtained pre-procedure and injury pathology assigned post-procedure. Synovial fluid was extracted and analyzed via liquid chromatography mass spectrometry metabolomic profiling to examine differences in metabolism between injury pathologies and participant sex. Additionally, samples were pooled and underwent fragmentation to identify metabolites.

**Results:** Metabolite profiles revealed that injury pathology phenotypes were distinct from each other where differences in endogenous repair pathways that are triggered post-injury were detected. Specifically, acute differences in metabolism mapped to amino acid metabolism, lipid-related oxidative metabolism, and inflammatory-associated pathways. Lastly, sexual dimorphic metabolic phenotypes were examined between male and female participants, and within injury pathology. Specifically, Cervonyl Carnitine and other identified metabolites differed in concentration between sexes.

**Conclusions:** The results of this study suggest that different injuries (e.g., ligament vs. meniscus), as well as sex are associated with distinct metabolic phenotypes. Considering these phenotypic associations, a greater understanding of metabolic mechanisms associated with specific injuries and PTOA development may yield data regarding how endogenous repair pathways differ between injury types. Furthermore, ongoing metabolomic analysis of synovial fluid in injured male and female patients can be performed to monitor PTOA development and progression.

**Clinical Relevance:** Extension of this work may potentially lead to the identification of biomarkers as well as drug targets that slow, stop, or reverse PTOA progression based on injury type and patient sex.

## 2. Introduction

Post-traumatic osteoarthritis (PTOA) accounts for approximately 12% of osteoarthritis (OA) cases equating to 5.6 million cases annually^1, 2^. One of the most prevalent risk factors that contributes to PTOA is joint injury. Within the United States, nearly 250,000 ACL injuries occur annually where approximately 70% of injured patients undergo ACL reconstruction^3, 4^. ACL injuries are frequently accompanied by damage to other tissues and structures within the knee such as the meniscus^5^. PTOA prevalence is influenced by the type of injury, where ACL injury alone has a prevalence range between 13-39%^2, 6–10^, whereas PTOA prevalence is significantly higher amongst those with combined ligament and meniscal injuries (21-48%)^2, 7, 9–13^.

Other patient-specific risk factors that contribute to PTOA include age, BMI, and sex. Annually, 15% of knee injuries were attributed to high-school athletes with young female athletes being twice as likely to sustain knee injuries requiring surgical repair compared to male athletes^14^. Furthermore, females are more likely to develop and experience more severe OA compared to males^15, 16^. These sex-differences can likely be attributed to hormonal and anatomical differences where females have wider pelvises, small femurs, thinner articular cartilage at the distal femur, a smaller ACL, and a narrower intercondylar notch^17–19^.

Although it is established that PTOA is associated with injury as well as other patient-specific risk factors, underlying mechanisms and metabolic alterations following injury at the joint level remain unknown. Examining acute differences in response to various injury types has the potential to positively influence patient treatment, outcomes, and reduce the burden of both PTOA and OA. The application of metabolomic profiling may identify biochemical phenotypes that represent and capture the physiological and metabolic status of the tissue of interest.

A handful of studies have used metabolomic profiling of synovial fluid (SF) to understand the pathology of OA^20–23^; however, the authors are not aware of any study using this method to quantitatively investigate acute metabolic perturbations induced by different knee injuries. By doing so, treatment and intervention may be improved to benefit overall patient health post-injury. Therefore, the primary goal was to examine metabolic alterations that differ between ligament (L), meniscal (M), and combined ligament and meniscal (LM) injuries to improve current understanding on the mechanisms triggered by acute injuries at the metabolic level.

The secondary goal of this study was to identify differences in pathway regulation and metabolite concentration between male and female participants. To accomplish both goals, metabolites were extracted from injured participants’ SF and analyzed via liquid chromatography-mass spectrometry (LC-MS). Global metabolomic profiling was applied to find specific metabolic perturbations associated with types of injury and patient sex. The identification of dysregulated metabolic pathways and metabolites may underpin mechanisms that differ between injury pathologies and those associated with PTOA. Moreover, ongoing metabolomic analysis of SF post-repair can be performed in conjunction with measurement of patient outcomes to oversee PTOA development and progression.

## 3. Materials & Methods

### a. Participant information and inclusion criteria

Under IRB approval, 58 participants were screened for eligibility between July 2021 and February 2022 at Virginia Commonwealth University. Participants were met in the pre-operative holding area on the day surgery and provided consent to participate in this study at that time. Inclusion criteria to participate in this cross-sectional study were (1) age between the age of 18-70 and (2) no history of prior knee injuries, chronic pain, or autoimmune disease(s). In total, SF was obtained from 45 knee arthroscopy patients (n=13 excluded) (Figure 1 and Supplementary Table 1). De-identified patient information provided included patient sex, age, BMI, and injury pathology. To limit potential bias, patient information including pathology, BMI, age, and sex were blinded throughout data analysis.

**Figure 1.**
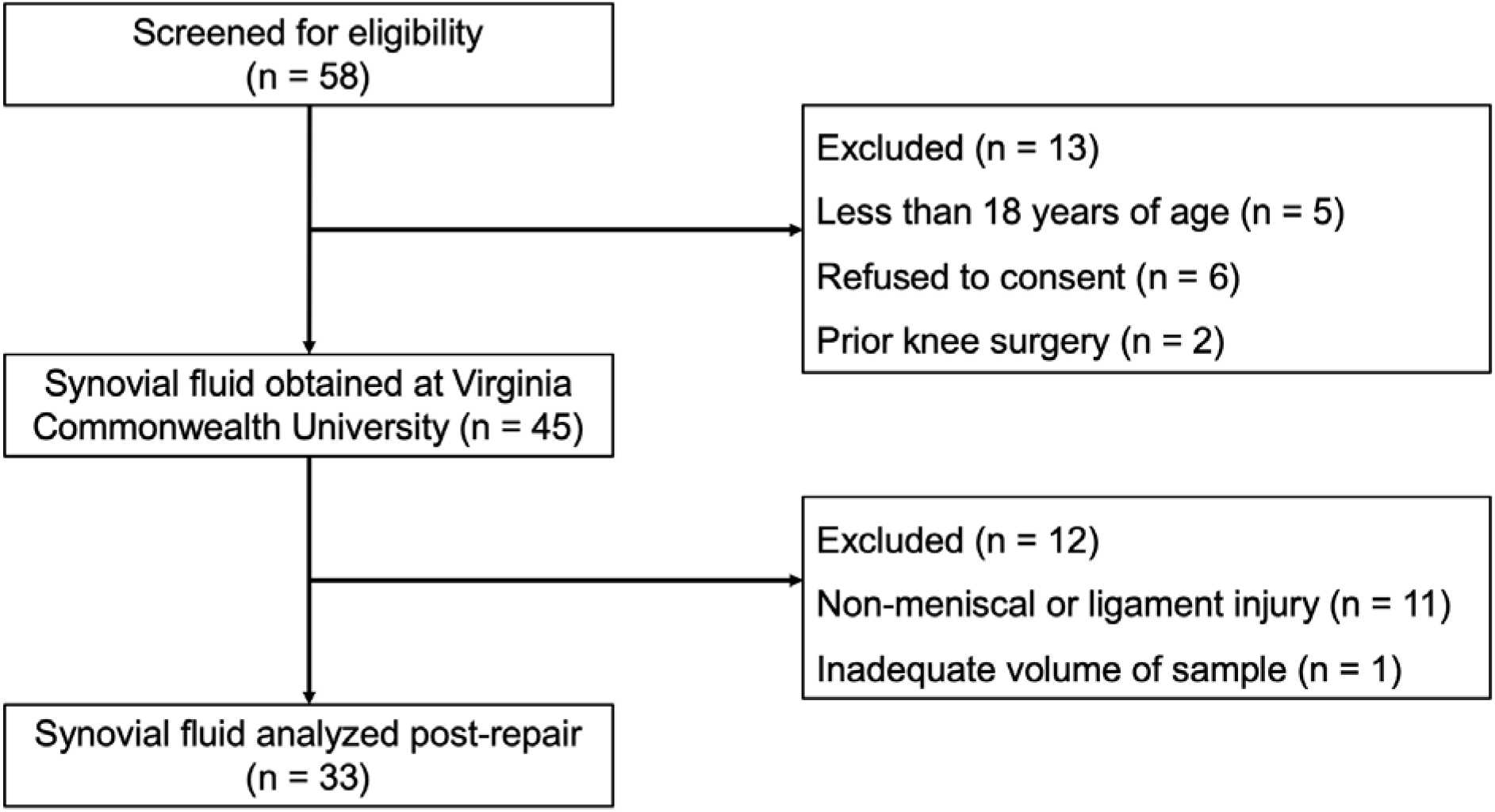
CONSORT Flow Diagram of the cross-sectional study.

### b. Synovial fluid sampling, extraction, and metabolic profiling

For all participants, SF was acquired in the operating room pre-procedure, and pathology assignment (i.e., right medial meniscus tear) was assigned post-procedure based on observations made in the operating room and the postoperative pathology report. Participant injury pathologies were categorized into one of three pathology groups: ligament (L), meniscal (M), and combined ligament and meniscal (LM) injuries (Table 1). SF from participants with non-meniscal or ligament injuries were excluded (n=13). In brief, all SF samples were extracted with methanol and acetonitrile, centrifuged, dried down via vacuum concentration, and prepped for LC-MS analysis. Two pooled samples were generated by randomly selecting 5 μL from extracted SF samples. All samples, participant SF and pooled samples were analyzed using a Waters Synapt XS. Full details on metabolite extraction and MS acquisition can be found in the supporting information,

**Table 1.** Participant information. Values indicate number of participants in each group.

| Sex | Ligament | Meniscus | Ligament & Meniscus |
| --- | --- | --- | --- |
| Female | 5 | 11 | 1 |
| Male | 5 | 7 | 4 |

Participant SF data were processed using MSConvert^24^ and XCMS^25^. MetaboAnalyst was utilized to statistically analyze samples, visualize dissimilarities, and pinpoint pathway dysregulation between males and females with different injuries. To identify metabolite features using MS^E^ data, Progenesis QI (Nonlinear Dynamics, Newcastle, UK) was utilized. Details about metabolomic profiling (statistical and pathway analyses) and identification can be found in the supporting information.

## 4. Results

### a. The synovial fluid metabolome differs by injury pathology

In total, 7,794 metabolite features were detected by LC-MS in the 33 SF samples. To assess global differences between injury pathologies, all metabolite features were analyzed using PCA and PLS-DA (Supplementary Fig. 1A-B). PCA displayed significant overlap of groups with PC1 and PC2 accounting for 44.6% of the variability in the dataset, whereas PLS-DA showed less overlap with Components 1 and 2 accounting for 33.3% of the variability in the dataset. Although changes at the global level were not detected, a group median heatmap analysis comparing the 25 highest VIP scoring metabolites was performed to examine and pinpoint phenotypic differences in regulation between pathology groups (Fig. 2A). Of the 25 features, 12 were more abundant in M injuries, whereas 13 of the 25 were more abundant in L injuries. Therefore, the data suggest injury type drives the regulation of the top 25 VIP Score metabolite features. Using MS^E^ data, analyses identified putative metabolites that differed in abundance between injury pathologies included Lysine, Tyrosine (L injuries); Glycerophosphocholine, Lycoperoside D (M injuries); Linoleic acid, Histidine (LM Injuries) and others (Fig. 2B, Supplementary Fig. 2, Supplementary Table 2). Therefore, when looking at all detected metabolite features, injury pathology does not change the global metabolic profile of SF. However, there are specific metabolic changes that are unique to each injury pathology as identified by PLS-DA VIP Scores and MS^E^ data.

**Figure 2.**
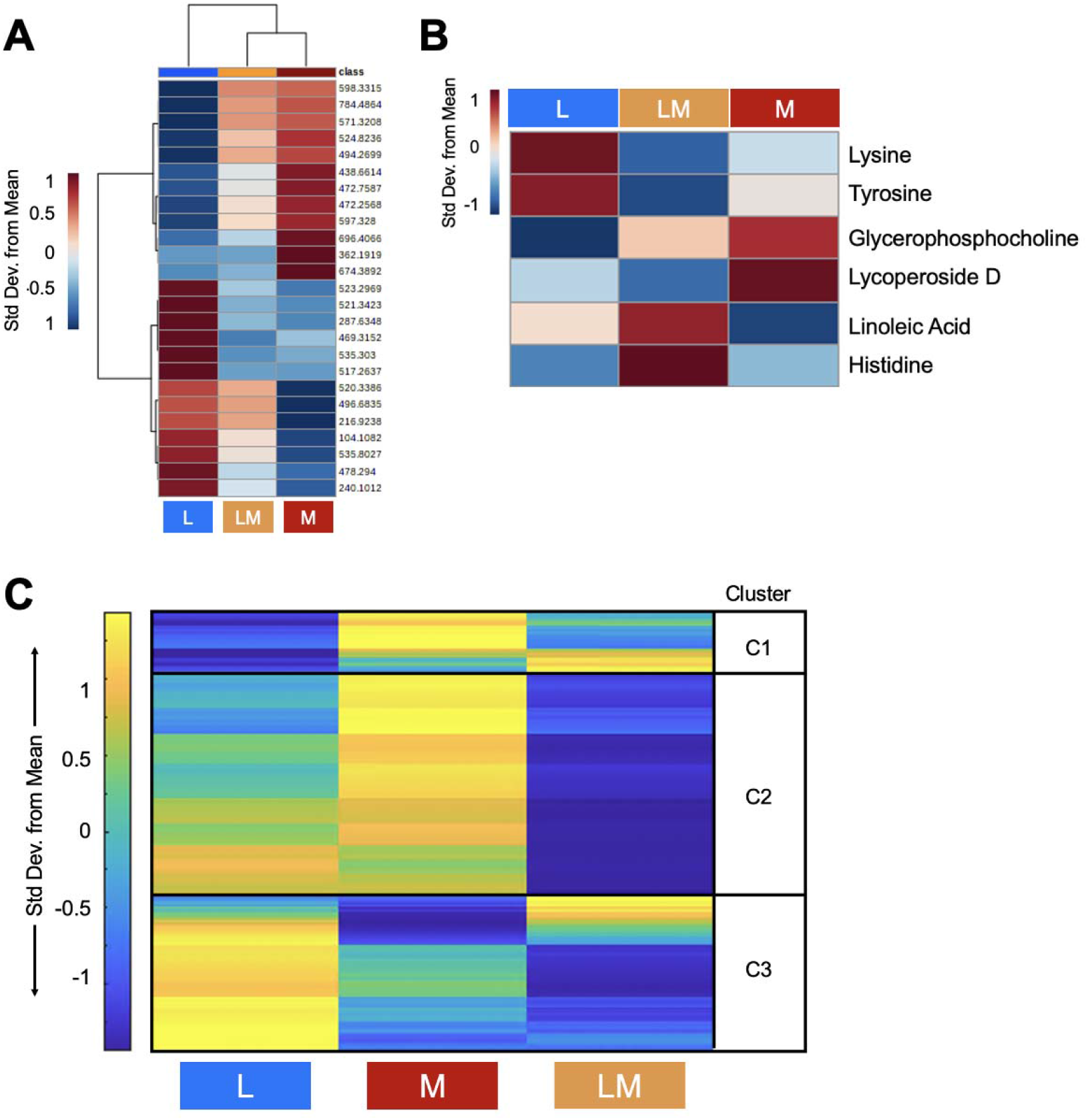
Metabolic phenotypes of participants that differ by injury pathology are distinct. (A) Plotting the top 25 PLS-DA Variable in Importance Projection Scores shows that different injury pathologies trigger distinct metabolic regulation patterns post-injury. Specifically, 12 of the 25 features are more abundant in meniscal injuries, whereas 13 of the 25 features were more abundant in L injuries. (B) MS/MS identified putative metabolites that differ in abundance between injury pathologies. Warmer colors (red) indicate higher abundance, while cooler colors (blue) indicate lower abundance. (C) Median metabolite intensity heatmap analysis of all metabolite features (7,994 distinct features) identified distinct metabolic phenotypes across injury pathologies and clusters of co-regulated metabolites. Clusters (1-3) were then subjected to pathway enrichment analyses to identify pathways that differed in regulation between injury pathologies. Warmer colors (yellow) and cooler colors (blue) indicate higher and lower metabolite intensities, respectively. The colors in A-C correspond to: Ligament injuries - light blue; Meniscal injuries - orange; Ligament and Meniscal injuries - red. L = ligament injuries. M = meniscal injuries. LM = ligament and meniscal injuries.

Functional pathway enrichment analyses were conducted to underpin endogenous repair pathways that differ between injury pathologies. To do so, a median metabolite intensity heatmap analysis was performed to visualize global changes across the metabolome to distinguish patterns, or clusters, of co-regulated and differentially expressed metabolite features (Fig. 2C). Clusters of metabolites identified underwent pathway enrichment analyses using MetaboAnalyst.

Metabolite features that had the highest concentration in L injuries, and the lowest in LM injuries, mapped to leukotriene metabolism, omega-6 fatty acid metabolism, vitamin E and B3 metabolism, mono-unsaturated fatty acid beta-oxidation, ubiquinone biosynthesis, butanoate metabolism, pyrimidine metabolism, and amino acid metabolism (arginine, proline, aspartate, asparagine, glycine, serine, alanine, threonine, tryptophan, lysine). Conversely, features that had the highest concentration in L injuries, and lowest in M injuries, mapped to C21-steroid hormone biosynthesis and metabolism, alkaloid biosynthesis, biopterin metabolism, glycerol- and glycosphingolipid metabolism, sialic acid metabolism, and glutathione metabolism. Metabolite features with the highest concentration in M injuries, and lowest in LM injuries, mapped to lipid-related beta oxidation pathways (di-unsaturated and saturated fatty acids beta-oxidation, dimethyl-branched-chain fatty acid mitochondrial beta-oxidation), the carnitine shuttle, and omega-3 fatty acid metabolism. Lastly, linoleate metabolism-associated metabolite features were the highest in LM injuries, and the lowest in M injuries (Table 2, Supplementary Table 3).

**Table 2.** Metabolic pathways associated with ligament, meniscal, and ligament and meniscal injuries identified by median metabolite intensity heatmap analysis. All reported pathways have a FDR-corrected significance level < 0.05. L = ligament injuries. M = meniscal injuries. LM = ligament and meniscal injuries. Clusters defined in Figure 2C.

| Cluster | Pathway | Highest | Lowest |
| --- | --- | --- | --- |
| 2 | Leukotriene metabolism | L | LM |
| 2 | Omega-6 fatty acid metabolism | L | LM |
| 2 | Vitamin E metabolism | L | LM |
| 2 | Arginine and Proline Metabolism | L | LM |
| 2 | Mono-unsaturated fatty acid beta-oxidation | L | LM |
| 2 | C5-Branched dibasic acid metabolism | L | LM |
| 2 | Ubiquinone Biosynthesis | L | LM |
| 2 | Vitamin B3 (nicotinate and nicotinamide) metabolism | L | LM |
| 3 | Aspartate and asparagine metabolism | L | LM |
| 3 | Glycine, serine, alanine and threonine metabolism | L | LM |
| 3 | Tryptophan metabolism | L | LM |
| 3 | Alanine and Aspartate Metabolism | L | LM |
| 3 | Butanoate metabolism | L | LM |
| 3 | Carnitine shuttle | L | LM |
| 3 | Lysine metabolism | L | LM |
| 3 | Pyrimidine metabolism | L | LM |
| 2 | C21-steroid hormone biosynthesis and metabolism | L | M |
| 2 | Alkaloid biosynthesis II | L | M |
| 2 | Biopterin metabolism | L | M |
| 3 | Glycerophospholipid metabolism | L | M |
| 3 | Glycosphingolipid metabolism | L | M |
| 3 | Histidine metabolism | L | M |
| 3 | Sialic acid metabolism | L | M |
| 3 | Glutathione Metabolism | L | M |
| 2 | Saturated fatty acids beta-oxidation | M | LM |
| 2 | Di-unsaturated fatty acid beta-oxidation | M | LM |
| 2 | Omega-3 fatty acid metabolism | M | LM |
| 2 | Dimethyl-branched-chain fatty acid mitochondrial beta-oxidation | M | LM |
| 2 | Linoleate metabolism | LM | M |

### b. Participant sex influences metabolomic profiles across injury pathologies

To determine if participant sex influences the SF metabolome, PLS-DA, fold change, and volcano plot analyses were conducted. PLS-DA revealed minimal overlap of male and female participants suggesting the SF metabolome reflects participant sex (Fig. 3A). Fold change analysis identified 262 metabolite features that were expressed in higher concentrations in males compared to females, and 24 metabolite features that were higher in females compared to males (Fig. 3B). Volcano plot analysis identified 17 metabolites features that were significant and had higher concentrations in males compared to females, and 11 metabolite features that were significant and in higher concentration in females compared to males (Fig. 3C).

**Figure 3.**
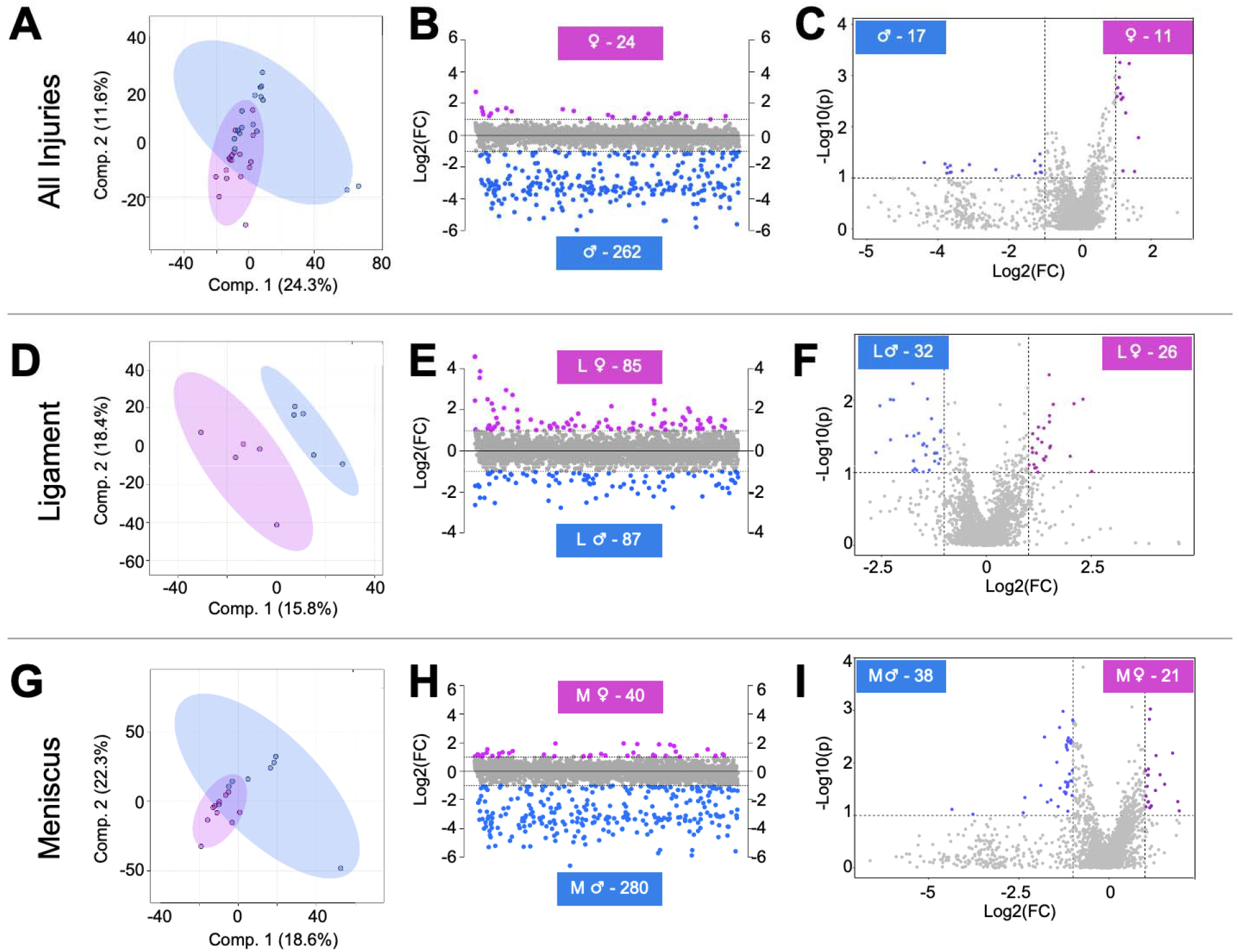
Metabolomic profiles of synovial fluid-derived metabolite features are associated with participant sex and injury pathology. (A) To assess metabolic differences associated with participant sex, all male and female participants not considering injury pathology, were included for PLS-DA analysis. This supervised statistical test revealed some overlap suggesting that the metabolome of male and female participants differ metabolically. (B) Fold change analysis identified 24 metabolite features with a FC > 2 in females, whereas 262 had a FC < −2 in males. (C) Volcano plot analysis identified 11 metabolite features in females that had a FC > 2 and p-value < 0.05. Conversely, 17 had a FC < −2 and a p-value < 0.05 in males. Similarly, (D) PLS-DA, (E) fold change, and (F) volcano plot analysis was applied to examine metabolic differences between male and female participants with ligament injuries and the same suite of analyses were applied to identify differences between male and female participants with meniscal injuries (G-I). Considering the identification of subpopulations of metabolite features that differ in regulation between male and female participants (A-C), male and female participants with ligament injuries (D-F), and male and female participants with meniscal injuries (G-I), it is evident that the metabolome is influenced by participant sex and injury pathology. The colors in A-I correspond to male and female participants: pink – females, blue – males. L = ligament injuries. M = meniscal injuries.

In a similar way, pairwise comparisons were performed to examine participant sex differences within injury pathology for L and M injuries. LM injuries were not examined due to insufficient sample size (LM Male n = 4, LM Female n = 1). Considering L injuries, PLS-DA displayed clear separation of participants that differ by sex (Fig. 3D). Fold change analysis identified 87 metabolite features that were expressed in higher concentrations in L males, whereas 85 were higher in concentration in L females (Fig. 3E). Volcano plot analysis identified 32 metabolite features that were significant and had higher concentrations in L males, whereas 26 were significant and in higher concentrations in L females (Fig. 3F). These same analyses were performed to analyze metabolic phenotypes associated with male and female M injuries. PLS-DA showed slight overlap of male and female participants with M injuries, further supporting the notion that differences at the metabolic differences are associated with participant sex (Fig. 3G). Fold change analysis identified 280 metabolite features that were highest in concentration in M males, whereas 40 features were highest in M Females (Fig. 3H). Volcano plot analysis identified 38 metabolite features that were significant and had higher concentration in M males, and 21 features that were significant and had higher concentrations in M females (Fig. 3I).

Populations of metabolite features from fold change and volcano plot analyses were then matched to identifications made using MS^E^ data. As expected with LC-MS, not all metabolite features were able to be identified (Supplementary Table 4). Cervonyl carnitine was detected in higher abundances in all females compared to males and when considering injury pathology (FDR corrected p-values < 0.05) (Fig. 4A-C). Phenylalanine was in higher abundances in all male participants compared to females. Considering sex and injury pathology, Cyclotricuspidoside C, Lucidenic acid N, and alpha-Chaconine were highest in males with M injuries compared to females with M injuries (Supplementary Figure 3A-D). Overall, these data strongly suggest that participant sex influences metabolic phenotypes across injury pathologies.

**Figure 4.**
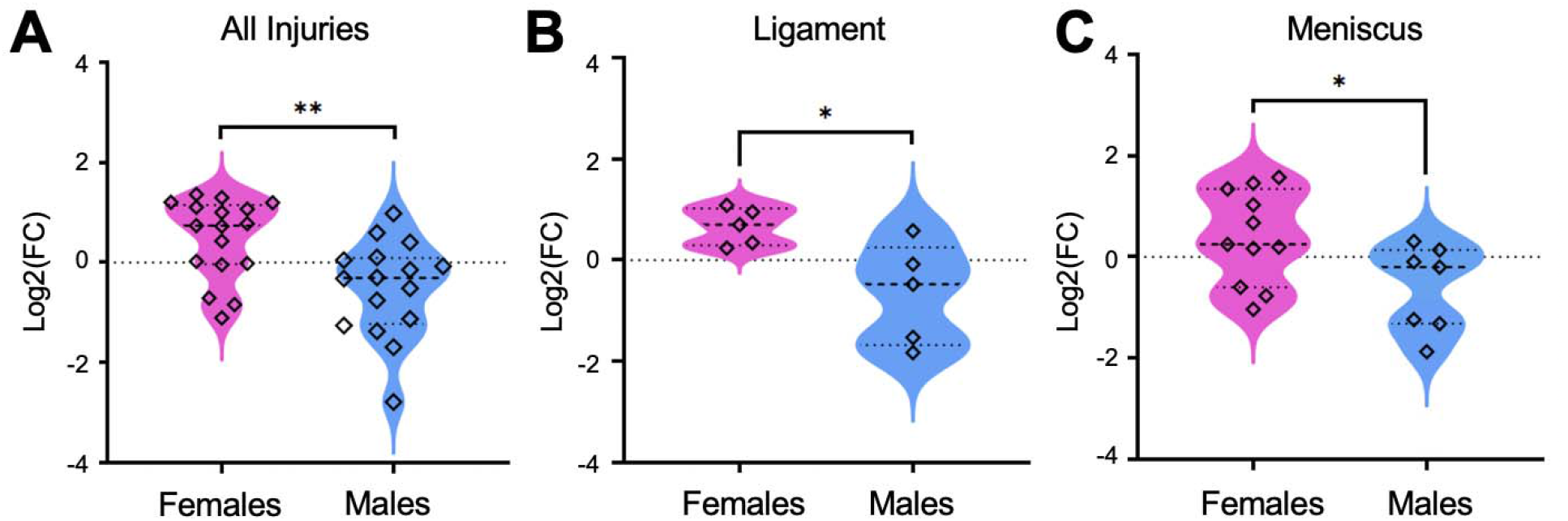
Cervonyl Carnitine differs in concentration between male and female participants. The identified metabolite, Cervonyl carnitine, was higher in concentration in (A) all female participants compared to males. Furthermore, this metabolite was higher in (B) females with ligament injuries and (C) meniscal injuries compared to males within the same injury pathology group. Data were initially evaluated using fold change and volcano plot analysis to pinpoint populations of metabolite features to identify, then features were matched to identified metabolites. Mass-to-charge intensities of interest were normalized and used to generate plots. To correct for multiple comparisons, FDR p-value corrections were performed and were less than < 0.05. Unpaired t-tests were performed in Graph Pad Prism. **Indicates p-value < 0.01 and *indicates p-value < 0.05. The colors correspond to male and female participants: pink – females, blue – males.

## 5. Discussion

To our knowledge, this is the first study to examine acute metabolic responses following different injury types in human SF. The goals of this study were to (1) identify metabolic perturbations that differ across injury pathologies including L, M, and combined LM injuries and (2)examine metabolic differences associated with sex. By applying LC-MS based metabolomics, metabolic phenotypes between injury pathologies vary suggesting different types of injury trigger specific acute metabolic responses. Metabolic phenotypes were associated with amino acid metabolism, lipid-related oxidative pathways, and various others. Additionally, SF-derived metabolites reflected participant sex, and male and female participants with similar injuries were metabolically distinct from each other. The detection of metabolites and generation of metabolomic phenotypes based on sex and injury pathology can be used to improve patient treatment and allow for more precise treatment to benefit joint and patient health post-injury.

### a. Acute metabolic responses post-injury varies based on injury pathology

Previous studies have applied metabolomics to examine metabolic shifts post-injury in animal models^26–29^, but no study to date has examined metabolic shifts in human SF induced by different injuries. Here, metabolic phenotypes were generated and differences in pathway regulation between injuries were detected. Specifically, amino acid metabolism (L injuries), lipid- and oxidative-related metabolism (M injuries), and inflammatory-associated pathways (LM injuries) differed in regulation between injury pathologies (Fig. 5).

**Figure 5.**
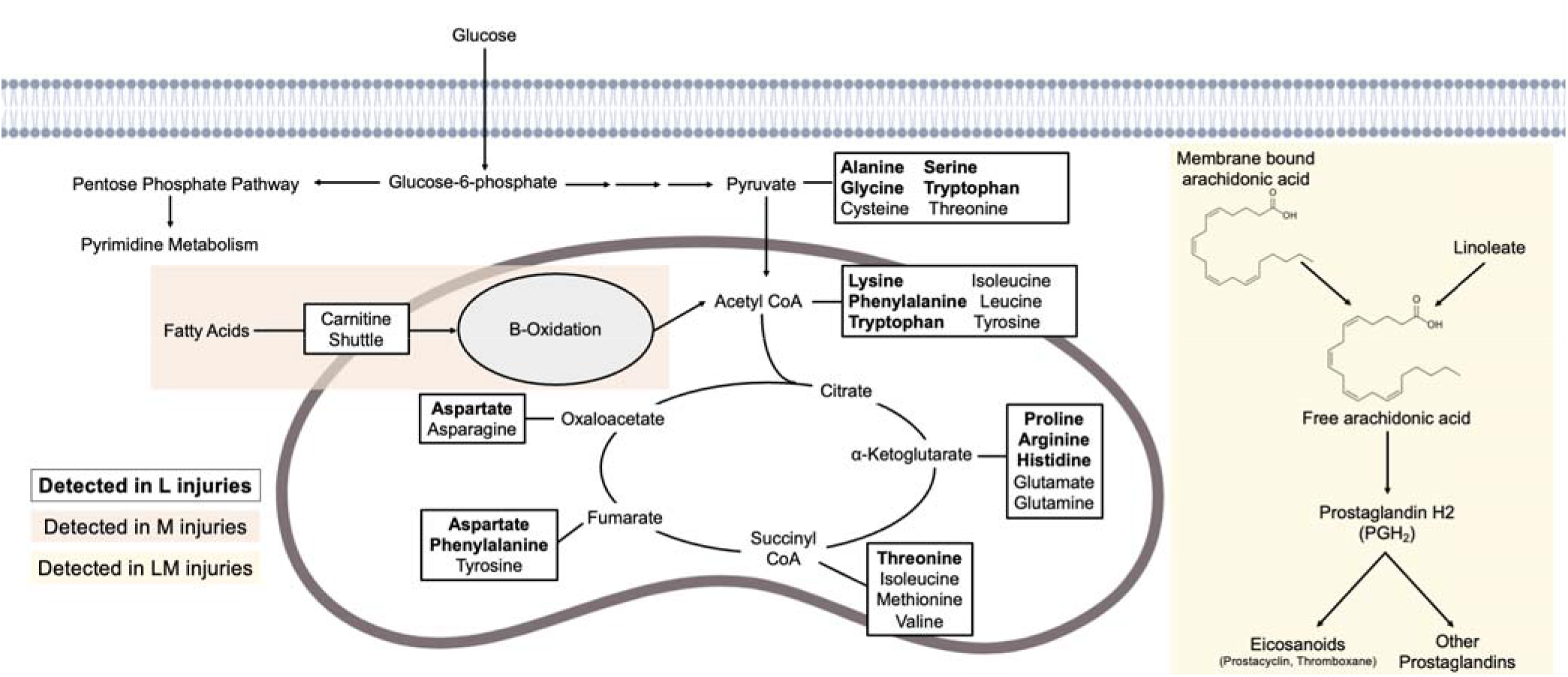
Alterations in synovial fluid metabolism post-injury are associated with different injury types. Metabolites that differ in abundance across ligament, meniscal, and ligament and meniscal injuries are associated with amino acid, lipid, and linoleate metabolism, respectively. Multiple amino acids, bolded, were detected in synovial fluid from participants with ligament injuries. Metabolites detected among meniscal injuries were related to lipid metabolism, specifically the carnitine shuttle and beta-oxidation (orange). Combined ligament and meniscal injuries were associated with linoleate metabolism, which is related to arachidonic acid metabolism and the generation of prostaglandins and eicosanoids (yellow).

Participants with L injuries had the greatest dysregulation of amino acid metabolism compared to M and LM injuries. Dysregulated amino acid pathways included asparagine, arginine, proline, lysine, and others. Asparagine and arginine, both having anti-inflammatory effects, have been detected in low concentrations in serum from OA patients.^30, 31^ Proline, a downstream product of arginine, contributes to collagen synthesis^32^ and has also been associated with OA^20, 31, 33, 34^. A prior study found higher concentrations of asparagine, arginine, and proline at day 1 that then decreased by day 8 in a non-invasive mouse ACL tear model^29^, suggesting these amino acids could be protective after injury and might be measured over time to monitor OA development. Similarly, NMR analysis of sheep SF following ACL injury found increased levels of serine and asparagine post-injury compared to non-injured sheep^28^. Using MS^E^ data, Lysine was identified and had the highest abundance in L SF. Lysine has previously been associated with collagen, the most abundant structural macromolecule of the extracellular matrix (ECM)^35^. Specifically, lysine residues undergo posttranslational modifications, catalyzed by lysl hydroxylases, to facilitate covalent cross-linking of collagen fibrils^36, 37^. These crosslinks are important as they strengthen collagen as well as supporting the assembly and stability of the ECM. Therefore, the identification and detection of Lysine in participants with ligament injuries may reflect a need for collagen synthesis post-injury. Combined, the magnified dysregulation of amino acid-related pathways in L participants, compared to M and LM participants, may reflect that L injuries have a higher cellular demand for amino acids for various reasons (i.e., energy generation, collagen synthesis, inflammation) compared to other injury pathologies. Few studies have determined the cellular density of healthy and injured human menisci and ligaments^38–40^, therefore, calculating cellular density in tandem with metabolomics could improve current understanding of metabolic changes relating to amino acid metabolism post-injury.

Amongst M participants, putatively identified metabolites related to lipid metabolism that were highest in abundance in M participants included Glycerophosphocholine and Lycoperoside D. Additionally, the carnitine shuttle and lipid-related oxidative metabolic pathways (di-unsaturated and saturated fatty acids β-oxidation, dimethyl-branched-chain fatty acid mitochondrial β-oxidation) were the most dysregulated compared to L and LM participants. The purpose of the carnitine shuttle is to transport fatty acids into the mitochondria to generate ATP via β-oxidation. Functioning mitochondria rely on glucose and fatty acids to yield energy, but dysfunctional mitochondria switch to relying on fatty acids more than glucose^41, 42^. This switch induced by various factors like cartilage breakdown during OA development can result in the accumulation of fatty acids^43, 44^ resulting in lower ATP production, impaired stress response, an increase in reactive oxygen species, apoptosis, and combined can lead to systemic irreparable damage^45–47^. Therefore, M injuries may trigger mitochondrial dysfunction leading to an increase in the shuttling of fatty acids into the mitochondria via carnitines where fatty acids become the primary energy source causing accumulation, mitochondrial dysfunction, and potentially, downstream PTOA or early-stage OA develop as a result. This response may be magnified in M injuries and suggests that distinct metabolic mechanisms are associated with specific injuries. Further examination of the metabolic regulation of carnitines and lipids as well as the role of metabolic dysfunction following injury may yield data on how differences in endogenous repair pathways contribute to PTOA development.

Lastly, linoleic acid, identified by MS^E^, had the highest abundance in LM SF. Additionally, linoleate metabolism was the most dysregulated pathway amongst LM participants compared to L and M participants. This pathway is upstream of arachidonic acid metabolism which generates fast-acting and short-lived signaling molecules such as the pro-inflammatory prostaglandin, PGH2, by cleaving and converting arachidonic acid^48–50^. Arachidonic acid has been detected in OA SF, specifically in higher levels in the early stages of OA and lower levels in late-stage OA^49, 51^. Therefore, the detection of linoleic acid and metabolites associated with linoleate metabolism in the present study may be associated with signaling and prostaglandin generation to regulate inflammation post-LM injuries and may serve as a marker to monitor PTOA development.

### b. The synovial fluid metabolome differs in association with participant sex and injury pathology

It is well established that knee injury, or trauma, is a risk factor for OA^1, 2^. Furthermore, females are at a higher risk for knee injury and downstream development of PTOA compared to males^14–16^. Although these empirical sex differences exist, diagnosis and treatment of males and females does not differ^52^. Therefore, the present detection of sex differences at the metabolic level build upon previous work that patient sex is a prevalent risk factor for PTOA. Specifically, metabolic phenotypes of SF-derived metabolite features were associated with participant sex and male and female participants with similar injury pathologies were metabolically distinct from each other. This is the first work, to our knowledge, to assess sex differences considering different injury pathologies in human SF.

Cervonyl carnitine was elevated in all female participants compared to all male participants. Carnitines have many systemic functions where they regulate oxidative and metabolic statuses, maintain membrane stability, and contribute to β-oxidation by transporting fatty acids into mitochondria^53–55^. Due to hormonal differences, oxidation of fatty acids is variable between males and females^56, 57^. Specifically, circulating estrogen levels are much higher in females than males, resulting in an increased expression of fatty acid oxidation proteins and pathways such as adenosine monophosphate-activated protein kinase (AMPK)^58, 59^. AMPK is a key energy sensor that maintains energy homeostasis, cellular metabolism, and promotes ATP production^60, 61^. The proposed mechanism for estrogen activation of AMPK entails estrogen binding to estrogen-receptor β (ERβ), causing an increase in Ca^2+^ stimulating Ca^2+^/Calmodulin-dependent protein kinase kinase β (CaMKKβ) to phosphorylate the AMPKα subunit where anabolism is inhibited and catabolism is activated resulting in the transport of fatty acids into the mitochondria to perform β-oxidation and generate ATP^62^ (Fig. 6). Atypical AMPK activity has been implicated in OA where mitochondrial function is impaired, ROS accumulate, and ATP production is reduced leading to cartilage degeneration, inflammation, and abnormal subchondral bone remodeling^61^. Therefore, the detection of elevated carnitine species in females compared to males may demonstrate that females rely on different metabolic pools and mechanisms, like AMPK activation via estrogen, to generate ATP and meet energy demands post-injury. Furthermore, carnitine-related species could be monitored overtime to gauge rates of β-oxidation, AMPK activity and function, joint health, and used to predict PTOA onset and development.

**Figure 6.**
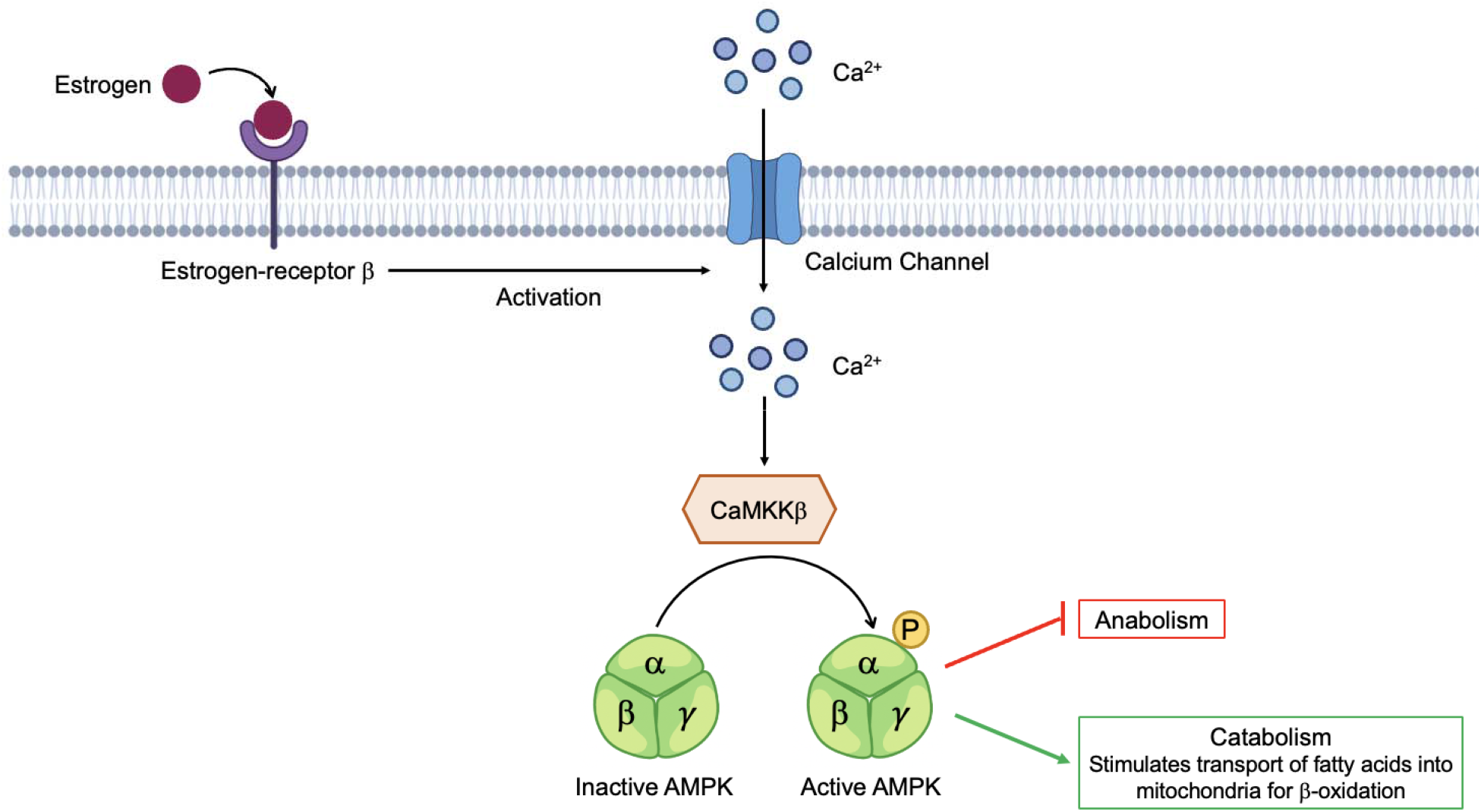
Proposed mechanism of AMPK activation by estrogen. Females have higher levels of circulating estrogen, which can bind estrogen-receptor β activating an influx of Ca^2+^ into the cell, stimulating Ca^2+^/Calmodulin-dependent protein kinase kinase β (CaMKKβ) phosphorylation of AMPK. At this point, anabolism is halted, and catabolism is stimulated resulting in the transport of fatty acids into the mitochondria, via carnitines, to perform β-oxidation. Differences in regulation of carnitine and lipid species between male and female participants may reflect reliance on different metabolic mechanisms, like estrogen activation of AMPK, and substrates for energy demands post-injury.

Taken together, these data suggest that participants that differ by sex and injury pathology have distinct metabolic phenotypes. The detection of carnitine-related species and others can be used to better understand metabolic mechanisms triggered by injury and how they vary between males and females. Further investigation of metabolite features dysregulated between males and females will improve patient treatment that accounts for sex considering females increased risk of ACL injury and OA development later in life.

### c. Limitations

While this study finds that metabolic phenotypes reflect patient sex and injury pathology, it is not without limitations. Firstly, participant SF was obtained by the same two surgeons in the operating room pre-procedure.However the time between injury and joint repair was not uniform across all participants. Therefore, varying windows of time without repair could have influenced metabolic results. Secondly, sample sizes for injury pathology groups were not uniform (M, n = 18; L, n = 10; LM, n = 5). Finally, the lack of healthy non-injured participants as well as participants with PTOA limits analyses and study conclusions.

### d. Conclusions

The results of this cross-sectional study demonstrate that injured males and females have distinct metabolic phenotypes. Considering sex-differences associated with rates of injury and PTOA development, the detection of dysregulated metabolites between male and females can be used to tailor patient treatment accordingly. Future studies aim to investigate the interaction between PTOA and sex, as well as other PTOA risk factors (i.e., BMI, age). Furthermore, this LC-MS based global approach provided clear differentiation between participants with L, M, and LM injuries. Specifically, differences in metabolism post-injury corresponded to amino acid (L injuries), lipid (M injuries), and linoleate metabolism (LM injuries). Considering these phenotypic associations, a greater understanding of metabolic mechanisms associated with specific injuries and PTOA may yield data regarding how endogenous repair pathways differ between injuries. Moreover, ongoing metabolomic analysis of SF in injured patients post-repair can be performed to monitor joint health as well as PTOA development. Completion of this work may potentially lead to the identification of biomarkers and drug targets that slow, stop, or reverse PTOA progression considering injury pathology and patient sex.

## Supporting information

STROBE Checklist

## 6. Acknowledgements

We thank Virginia Commonwealth University as well as the Department of Orthopaedic Surgery for assisting in obtaining participant synovial fluid used in this cross-sectional study. Additionally, we thank the Mass Spectrometry Facility at Montana State University including Dr. Donald Smith and Jesse Thomas for assisting in LC-MS analysis, interpretation, and metabolite identification. Funding for the Mass Spectrometry Facility used in this publication was made possible by the M.J. Murdock Charitable Trust, the National Institute of General Medical Sciences of the National Institutes of Health (P20GM103474 and S10OD28650). This study was funded by the National Science Foundation under Award Number CMMI 1554708 (RKJ), and the National Institutes of Health under Award Numbers R01AR073964 (RKJ).

## Author Contributions

HDW designed experiments, performed metabolite extractions, ran LC-MS samples, analyzed data, and drafted manuscript. AHW performed metabolite extractions and analyzed data. PP, JS, RO, and ARV obtained samples and provided clinical input. AV and RO performed repairs, and assisted in designing experiments. BB and RKJ designed experiments and analyzed data. All authors have read and revised the manuscript.

---

**Supplementary Table 1**. Participant information for all participants including BMI, age, sex, and reason for knee arthroscopy repair.

**Supplementary Table 2**. Identified putative metabolites that differ in abundance between injury pathology identified by MS^E^ with a score > 60 and fragmentation score > 12.

**Supplementary Table 3**. All metabolic pathways determined from MetaboAnalyst when comparing participants with different injury pathologies (Ligament, Meniscal, Ligament and Meniscal injuries) using median metabolite intensity heatmap analysis. Clusters defined on Figure 2C.

**Supplementary Table 4**. Identified putative metabolites that differ in abundance between male and female participants identified by MS^E^ with a score > 60 and fragmentation score > 12.

**Supplementary Figure 1.**
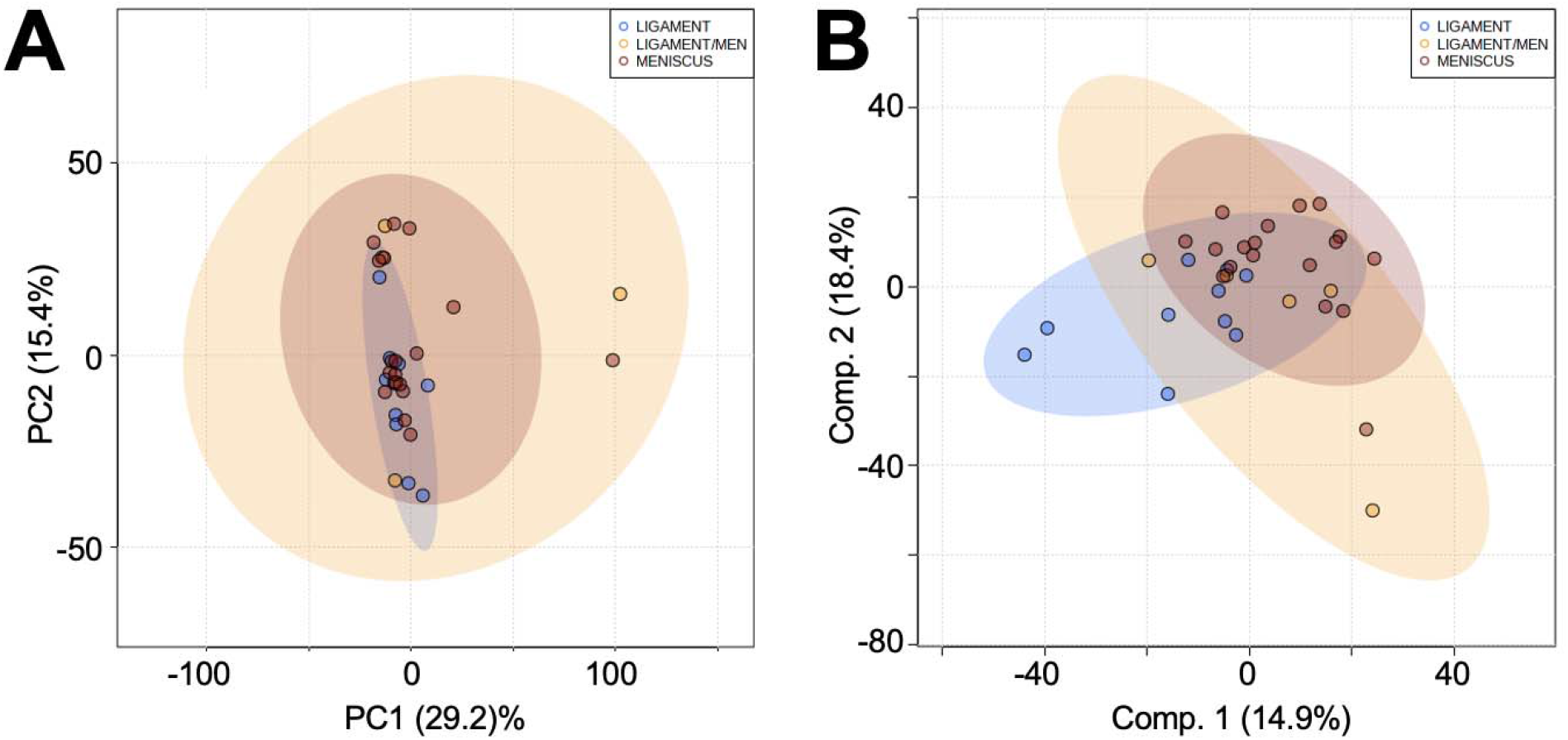
Global profiles considering all detected metabolite features are similar across injury pathologies. Unsupervised and supervised multivariate analyses display overlap in the global profiles of different injury pathologies. (A) PCA, an unsupervised test, shows significant overlap when considering all 7,794 features. PC1 and PC2 combined account for 44.6% of the variability in the dataset. (B) PLS-DA, a supervised test, shows less overlap of groups. Components 1 and 2 account for 33.3% of the variability of the dataset. These two tests combined suggest that global profiles generated by all metabolite features detected do not differ and that additional analyses are required to pinpoint specific phenotypic changes. The colors in A and B correspond to: Ligament injuries - light blue; Meniscal injuries - orange; Ligament and Meniscal injuries - red. L = ligament injuries. M = meniscal injuries. LM = ligament and meniscal injuries.

**Supplementary Figure 2.**
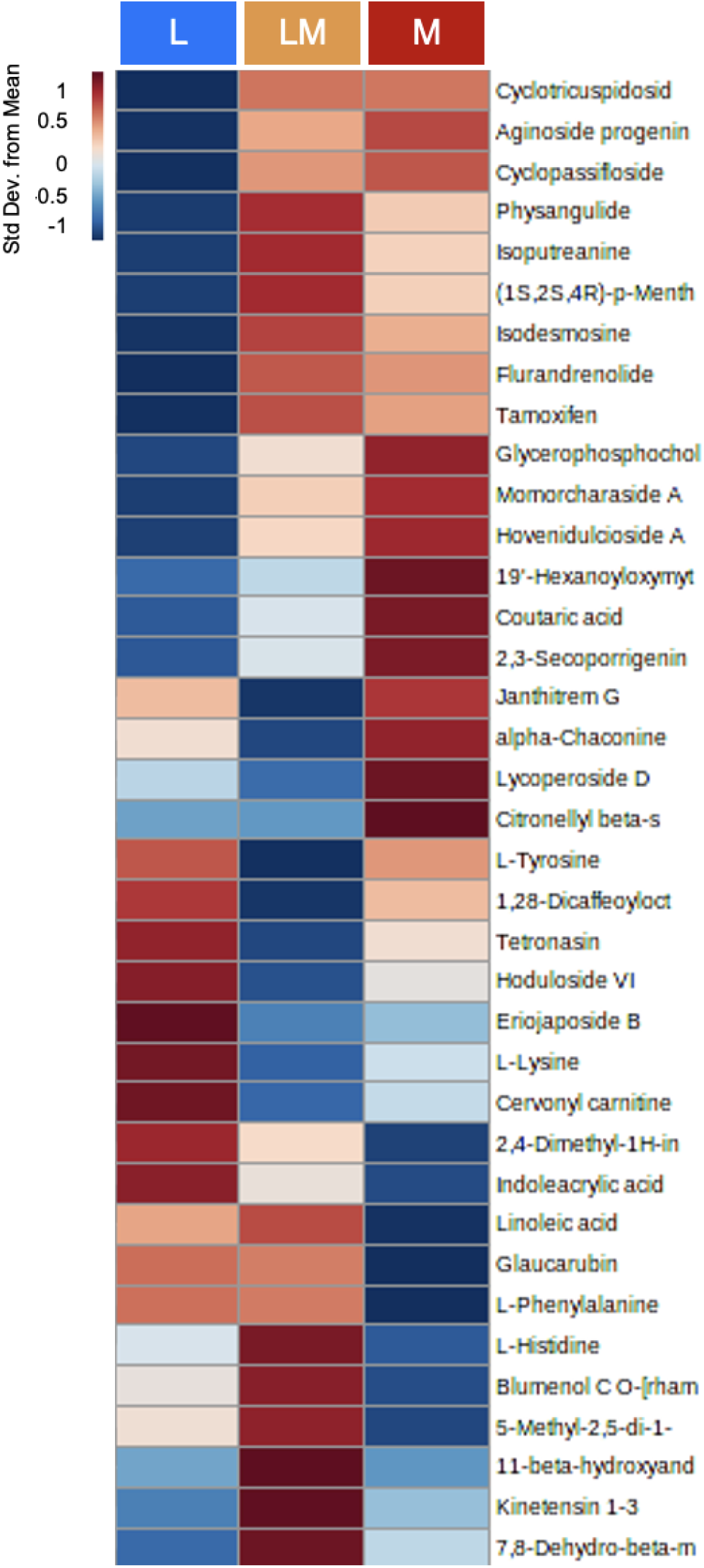
All putative metabolite identifications visualized by a heatmap. Warmer colors (red) indicate higher abundance, while cooler colors (blue) indicate lower abundance.

**Supplementary Figure 3.**
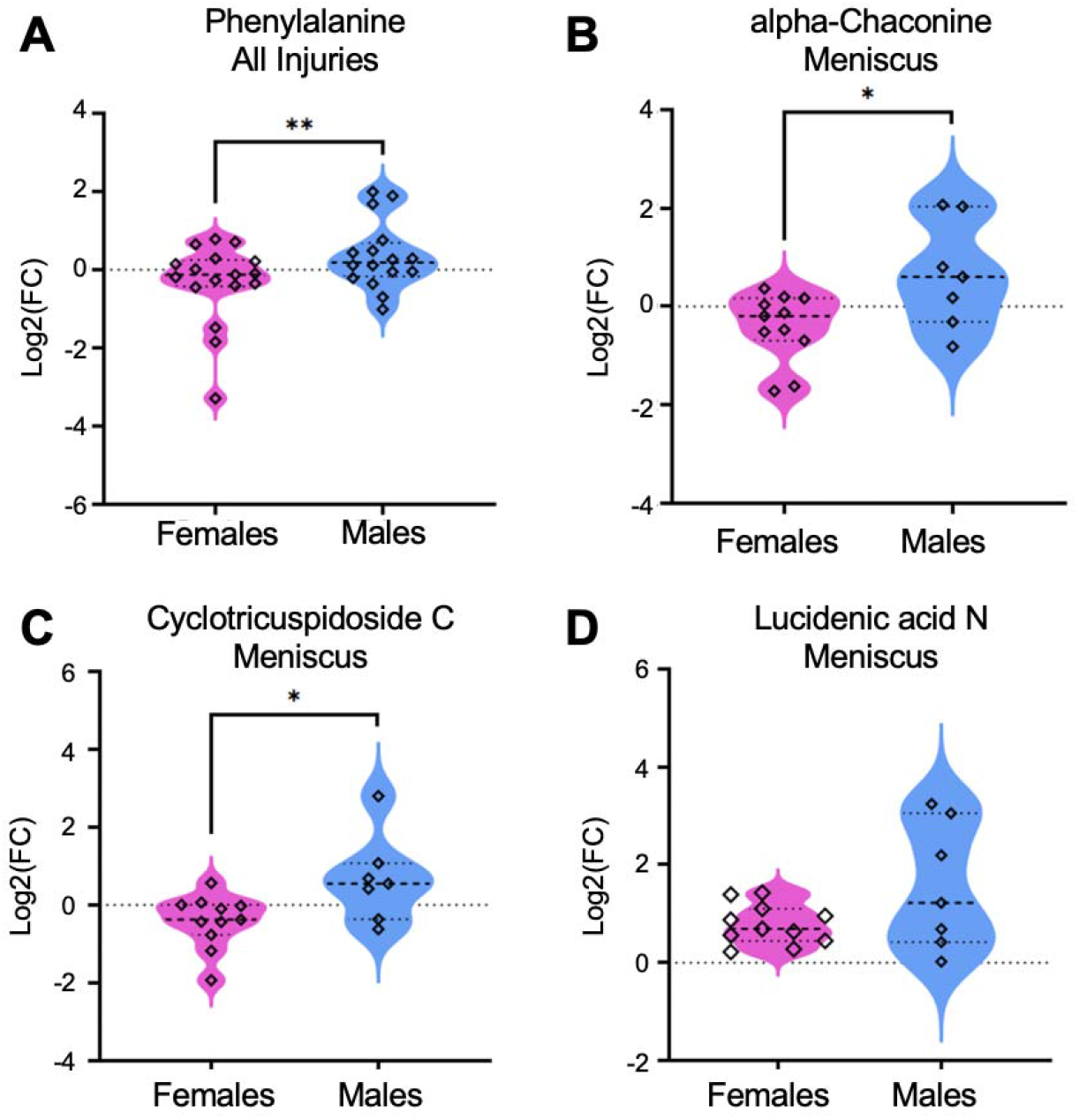
Concentration of identified metabolites differ in concentration between male and female participants and those with meniscal injuries. Using fold change and volcano plot analyses and MS^E^ data, populations of metabolites deemed significant were searched against identified metabolites. Of these, (A) Phenylalanine was in higher abundances in all male participants compared to females. (B) Alpha-Chaconine, (C) Cyclotricuspidoside C, and (D) Lucidenic acid N were highest in abundance in males with meniscal injuries compared to females with this same injury. Mass-to-charge intensities of interest were normalized and used to generate plots. To correct for multiple comparisons, FDR p-value corrections were performed and were less than < 0.05. Unpaired t-tests were performed in Graph Pad Prism. **Indicates p-value < 0.01 and *indicates p-value < 0.05. The colors in A-B correspond to male and female participants: pink – females, blue – males.

