## Supporting Information for "Metabolomic Phenotypes Reflect Patient Sex and Injury Status: A Cross-Sectional Analysis of Human Synovial Fluid"

Additional File 1: Supplementary information for publication

### 1.1 Synovial Fluid metabolite extraction protocol

In total, 100 µL of participant SF samples were aliquoted, centrifuged at 500 x g for 5 minutes at 4°C to remove cells and debris, and precipitated with 500 µL of 4:1 methanol:$H_2O$. Samples were vortexed for 1 minute then stored at -20°C for 5 minutes. This process was repeated 5 times, then samples were stored at -20°C overnight. The following day, samples were pelleted by centrifugation at 16,100 x g for 10 minutes at 4°C. Supernatants were harvested and dried via vacuum concentration to evaporate solvent and isolate metabolites. To further extract SF metabolites, dried samples were resuspended with 250 µL of 1:1 acetonitrile:water. Again, precipitated SF samples were vortexed for 1 minute then stored at -20°C for 5 minutes. Next, samples were centrifuged at 16,100 x g for 10 minutes at 4°C and supernatants were transferred to new Eppendorf vials for vacuum concentration. The remaining dried samples were resuspended with 50 µL of 1:1 acetonitrile:water for LC-MS analysis. At this time, two pooled samples were generated by randomly selecting 5 µL of extracted SF from 10 participants. Two quality control samples were created, one containing only mass spectrometry grade water and another that underwent metabolite extraction using only the solvents.

### 1.2 Liquid chromatography-mass spectrometry

All samples, participant SF, pooled, and quality control samples, were analyzed using an Acquity UPLC plus coupled through an electrospray ionization source to a Waters Synapt XS. Metabolites were separated using a Cogent Diamond Hydride HILIC column (150 x 2.1 mm) at a flow rate of 0.400 µL/min. Solvents used were 95% water 5% acetonitrile with 0.1% formic acid (solvent A) and 95% acetonitrile 5% water with 0.1% formic acid (solvent B). The 19-minute elution gradient consisted of 95 to 25% solvent B over 12 minutes, and each run began with 2 minutes of wash. Quality control blanks were injected periodically throughout the overall run to minimize spectral drift and assess LC-MS performance. Participant SF samples underwent standard MS1. Data independent acquisition, $MS^E$, was performed on pooled samples with a constant high energy ramp of 30-50V.

### 1.3 Global metabolomic profiling: statistics and pathway enrichment analyses

LC-MS data consisting of mass-to-charge ratios (m/z values), relative metabolite abundance, and retention times were processed using MSConvert[24] and XCMS[25]. To correct for non-normal distributions, all data underwent interquartile range normalization, were log-transformed, and autoscaled (mean-centered and divided by standard deviation of each variable).

To statistically analyze samples and visualize dissimilarities in metabolomic profiles between males and females with different injuries, MetaboAnalyst was used[26]. Specifically, principal component analysis (PCA) and partial least squares-discriminant analysis (PLS-DA) were used to visualize the presence, or absence, of overlap between participant derived-SF metabolic phenotypes. Variable Importance in Projection (VIP) Scores, an extension of PLS-DA where metabolite features are scored based on their contribution to discriminating between cohorts, were also considered to better understand metabolic shifts induced by different injury types. Volcano plot, fold change, and heatmap analyses were performed to analyze individual(s), and clusters, of co-regulated metabolite features for pathway enrichment analyses using MetaboAnalyst's Functional Analysis feature. This module employs the *mummichog* algorithm which analyzes the inputted metabolite features to predict networks of functional activity and derive biological relevance. Pathway library Human MFN was reference to match metabolite features to putatively identified metabolites (mass tolerance: 5 ppm, positive mode). Significance was determined using an FDR-corrected significance level of $p < 0.05$ decided on *a priori*.

To identify metabolite features using $MS^E$ data, Progenesis QI (Nonlinear Dynamics, Newcastle, UK) was utilized. All MS1 and MS2 centroided data was imported, peak picked, and aligned. The Human Metabolome Database (HMDB) was utilized to compare acquired and theoretical fragmentation for identification purposes. Identifications found were then crosschecked by retention time and fragmentation score. For a metabolite identity to be accepted, metabolites must receive a Progenesis score greater than 60/100 and a fragmentation score greater than 12. The three properties that contributed to these scores and percentages were mass error, isotope distribution similarity, and retention time error. Parts per million (ppm) error was calculated between MS1 and $MS^E$ data, and identifications with error > 20 ppm were excluded. Identities made were compared to metabolite features of interest. To correct for multiple comparisons, FDR p-value corrections for identified metabolites were performed.
